# Removal of rare amplicon sequence variants from 16S rRNA gene sequence surveys biases the interpretation of community structure data

**DOI:** 10.1101/2020.12.11.422279

**Authors:** Patrick D. Schloss

## Abstract

Methods for remediating PCR and sequencing artifacts in 16S rRNA gene sequence collections are in continuous development and have significant ramifications on the inferences that can be drawn. A common approach is to remove rare amplcon sequence variants (ASVs) from datasets. But, the definition of rarity is generally selected without regard for the number of sequences in the samples or the variation in sequencing depth across samples within a study. I analyzed the impact of removing rare ASVs on metrics of alpha and beta diversity using samples collected across 12 published datasets. Removal of rare ASVs significantly decreased the number of ASVs and operational taxonomic units as well as their diversity. Furthermore, their removal increased the variation in community structure between samples. When simulating a known effect size, removal of rare ASVs reduced the power to detect the effect relative to not removing rare ASVs. Removal of rare ASVs did not affect the false detection rate when samples were randomized to simulate a null model. However, the false detection rate increased when rare ASVs were removed using a null distribution and assignment of samples to simulated treatment groups according to their sequencing depth. The false detection rate did not vary when rare ASVs were retained. This analysis demonstrates the problems inherent in removing rare ASVs. Researchers are encouraged to retain rare ASVs, to select approaches that minimize PCR and sequencing artifacts, and to use rarefaction to control for uneven sequencing effort.

**Importance:** Removing rare amplicon sequence variants (ASVs) from 16S rRNA gene sequence collections is an approach that has grown in popularity for limiting PCR and sequencing artifacts. Yet, it is unclear what impact an abundance-based filter has on downstream analyses. To investigate the effects of removing rare ASVs, I analyzed the community distributions found in the samples of 12 published datasets. Analysis of these data and simulations based on them showed that removal of rare ASVs distorts the representation of microbial communities. This has the effect of artificially making it more difficult to detect differences between treatment groups. Also of concern was the observation that if sequencing depth is confounded with the treatment, then the probability of falsely detecting a difference between the treatment groups increased with the removal of rare ASVs. The practice of removing rare ASVs should stop, lest researcher adversely affect the interpretation of their data.

## Introduction

16S rRNA gene sequencing is a mainstay of microbial community analysis (1). Two elements that are held in tension in the analysis of 16S rRNA gene sequence data are how to adequately remove PCR and sequencing artifacts while decreasing the granularity of the taxonomic level that is used in the analysis. When coarse taxonomic levels (e.g. genus level) are used, the effects of artifacts are minimized since the genetic breadth of the level is wider than the diversity of artifacts. Conversely, with fine taxonomic levels (e.g. amplicon sequence variants; ASVs) the effects of artifacts are significant since each artifact may represent a new ASV.

Numerous studies have attempted to address the problem of removing or “denoising” artifacts from data generated using Illumina’s MiSeq platform. In one approach, paired sequence reads are aligned and any discrepencies between the reads are resolved based on the difference in quality score for the position in question (2–4); quality scores are also used to curate single reads (5). In addition, a polishing step is often used to identify ASVs based on the frequency and similarity of sequences (2, 3, 6). In a second approach, the quality scores and types of errors are modelled to cluster sequence reads directly into ASVs (7). Regardless of the approach, many pipelines advocate for abundance-based screening where rare sequences are removed from each dataset prior to outputting the sequence data as ASVs (3, 6, 7). Some algorithms recommend removing all ASVs that appear one (i.e. singletons) (7), eight (3), or ten (6) or fewer times; these pipelines also vary in whether the minimum abundance threshold should be applied to individual samples (6, 7) or the pool of samples in a study (3). A notable exception, the mothur-based pipeline discourages the practice of removing rare sequences (2). After assigning sequences to ASVs, ASVs are often analyzed as a taxonomic unit or clustered to generate operational taxonomic units (OTUs) or phylotypes.

The abundance-based screening approach assumes that rare ASVs are more likely to be artifacts than more abundant ASVs. Sequencing of mock communities confirms that artifacts tend to be rare (2, 5). Proponents of abundance-based screening point to their ability to obtain the correct number of ASVs, OTUs, or phylotypes with data generated from sequencing mock communities when rare ASVs are removed (5). However, this approach effectively overfits the curation pipeline to data generated from a phylogenetically simple community with an atypical community distribution that is often sequenced to a coverage that is not achieved with biological samples. It is necessary to think more deeply about the practice of abundance-based screening.

The minimum abundance thresholds that have been proscribed were developed and applied without regard for the number of sequences generated from each sample. These recommendations appear to assume that the number of reads per sample is consistent within and between studies. However, it is common that the number of sequences generated from each sample may vary by two or three orders of magnitude (e.g. Table 1 and Figure S1). An ASV that appears once in a sample with 2,000 sequences is more trustworthy than an ASV that appears once in a sample with 100,000 sequences since it has a 50-fold higher relative abundance. But, according to the pipeline recommendations, they are treated as being equally trustworthy. Rather than removing rare ASVs, the approach taken by the mothur pipeline applies the classical ecological approach of rarefaction. Each sample is rarefied to the same sequencing depth so that the number of artifacts that appears in each sample is controlled.

**Table 1.** Summary of studies used in the analysis. For all studies, the number of sequences used from each study was rarefied to the smallest sample size. A graphical represenation of the distribution of sample sizes for each study and the samples that were removed from each study are provided in Figure S1.

| Study (Ref) | Samples | Total sequences | Median sequences | Range of sequences | Fold-difference between largest and smallest sample | SRA study accession |
| --- | --- | --- | --- | --- | --- | --- |
| Bioethanol (22) | 95 | 3,972,943 | 16,015 | 3,688-356,136 | 96.6 | SRP055545 |
| Human (23) | 490 | 20,909,768 | 32,505 | 10,523-430,415 | 40.9 | SRP062005 |
| Lake (24) | 52 | 3,169,868 | 69,041 | 15,347-112,871 | 7.4 | SRP050963 |
| Marine (25) | 7 | 1,391,396 | 193,464 | 133,516-254,060 | 1.9 | SRP068101 |
| Mice (2) | 348 | 2,813,747 | 6,477 | 1,804-30,565 | 16.9 | SRP192323 |
| Peromyscus (26) | 111 | 1,555,545 | 12,446 | 4,464-33,644 | 7.5 | SRP044050 |
| Rainforest (27) | 69 | 946,295 | 11,561 | 4,932-37,767 | 7.7 | ERP023747 |
| Rice (28) | 490 | 22,591,168 | 43,216 | 2,776-193,464 | 69.7 | SRP044745 |
| Seagrass (29) | 286 | 4,130,454 | 13,567 | 1,803-45,191 | 25.1 | SRP092441 |
| Sediment (30) | 58 | 1,154,174 | 17,584 | 7,685-68,321 | 8.9 | SRP097192 |
| Soil (31) | 18 | 956,656 | 51,844 | 47,806-59,956 | 1.3 | ERP012016 |
| Stream (32) | 201 | 21,162,574 | 90,159 | 9,175-390,964 | 42.6 | SRP075852 |

Experience sequencing biological samples also demonstrates that there are bona fide ASVs that may have an abundance below the proscribed threshold. For example, the abundance of an ASV may be below the threshold in some samples or time points and above the threshold in others. However, rarity, both in terms of prevalence and incidence, is an important ecological concept (8). Removing rare ASVs likely hinders one’s to ability to make inferences about the dynamics and nature of the populations that rare ASVs represent. Furthermore, removing ASVs whose abundances are below the proscribed threshold also potentially biases the community structure of the samples.

In the current study, I used published sequence data from 12 studies to investigate the nature of rare ASVs (i.e. those that appear 10 or fewer times) and the effect that removing them has on downstream analysis of microbial communities. The analysis was also performed using traditional OTUs, where ASVs subjected to abundance-based screening were clustered such that the ASVs within an OTU were no more than 3% different from each other. The results reject the assumptions built into abundance-based screening and highlight the problems inherent in removing rare ASVs.

## Results

### Datasets

I collected 12 publicly available datasets that used the Illumina MiSeq platform to sequence the V4 region of the 16S rRNA gene from a variety of environments (Table 1; Figure S1). After removing poor quality and chimeric ASVs and samples that had uncharacteristically low number of sequences for the dataset, these datasets included between 7 and 490 samples. The median number of sequences for each dataset ranged between 6,477 and 193,464. Strikingly, aside from the relatively small marine and soil datasets, the difference between the sample with the fewest sequences and the sample with the most sequences for each dataset varied by between 7.4 and 96.6-fold.

### The nature of singletons

Removal of rare ASVs is commonly justified as a method of removing ASVs that are artifacts. If such ASVs are artifacts, then one would expect the number of singleton ASVs to accumulate with sequencing depth. Contrary to this expectation, the median percentage of sequences that were discarded when singleton ASVs were removed from each dataset varied between 0.42 and 22.23% (bioethanol and seagrass). In addition, with the exception of the samples from the marine and sediment datasets (Spearman correlation, P>0.05), the fraction of singleton ASVs in samples was negatively correlated with the number of sequences in each sample with a range between −0.27 and −0.87 (rice and bioethanol) (Figure 1A). This showed that with additional sequencing, the probability of seeing singleton ASVs in multiple samples was greater than the probability of generating an artifact. This suggests that the singleton ASVs are not as likely to be artifacts as previously thought. Furthermore, if singleton ASVs were artifacts, then one would not expect to find them in other samples from the same dataset. In fact, singleton ASVs from samples with fewer sequences were often found in samples with more sequences. At least 50% of the singleton ASVs found in the samples from the mice, rice, seagrass, and stream datasets were found in another sample from the same dataset (Figure 1B). Considering the likelihood of finding an ASV duplicated in another sample is confounded by the number of samples and inter-sample diversity, the high coverage of singleton ASVs in these datasets was remarkable. The correlation between the number of sequences in a sample and the fraction of that sample’s singleton ASVs that were covered by another sample in the dataset was significant and negative for 9 of the datasets ranging between −0.31 and −0.84 for the rice and seagrass datasets, respectively (Figure 1C). The negative correlation indicated that the singleton ASVs in the smaller samples were more likely to be covered by ASVs in the larger samples. Among the three datasets without a significant correlation (Spearman correlation, P>0.05), the marine and soil datasets had the fewest samples in our collection and the stream dataset already had a high level of coverage regardless of the number of sequences. Contrary to the common motivation for removing rare ASVs, these results indicate that this practice disproportionately impacts samples with fewer sequences and likely removes more non-artifact ASVs than those that are artifacts.

**Figure 1.**
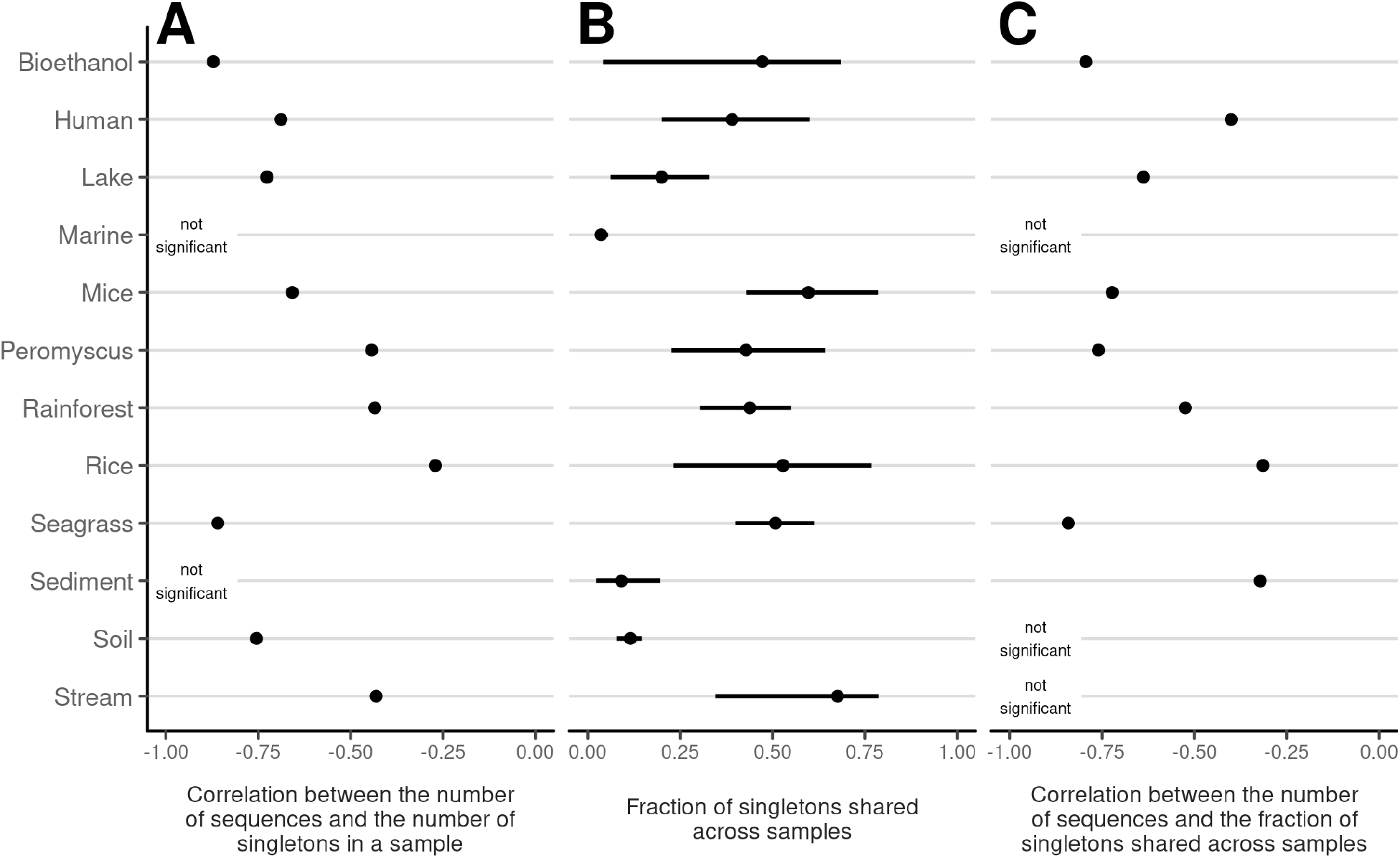
Singletons are more common in samples with fewer seqeunces and tend to be shared with samples having more sequences. For each of the 12 datasets, Spearman correlation coefficients were calcualted between the number of sequences in each sample and the number of singletons in the sample (A) and the fraction of its singletons that were shared with another sample (C). Those correlations that were not statisically significant had a P-value greater than 0.05. The faction of singletons shared across samples (B) were calculated for each dataset. The median value is shown with a solid circle and the 95% confidence interval is indicated by the solid line.

### The impact of removing rare ASVs on the information represented in each sample

Removing rare ASVs will reduce the richness of ASVs (i.e. the number of ASVs per sample) and increase the relative abundance of the remaining ASVs. To quantify the effect of removing rare ASVs on the information contained within each sample, I varied the minimum abundance threshold to simulate removing ASVs of varying rarity from each sample. The richness of ASVs in each sample decreased by between 34.4 and 86.2% (peromyscus and soil) when removing those ASVs that only appeared once and by between 76.0 and 95.6% (sediment and soil) when removing those that appeared ten or fewer times from each sample (Figure 2A). Similarly, the Shannon diversity decreased by between 1.8 and 15.9% (human and soil) when removing ASVs that only appeared once and by between 5.4 and 35.4% (human and seagrass) when removing ASVs that appeared ten or fewer times from each sample (Figure 2B). Next, I assigned the ASVs to OTUs, which were defined as a group of ASVs that were more than 97% similar to each other to assess the impact of removing rare ASVs on higher level taxonomic groupings that are commonly used in microbial ecology studies. Although pooling similar ASVs into OTUs reduced the impact of removing the rare ASVs relative to the ASV-based analysis, the minimum abundance threshold still decreased the richness of OTUs and the diversity decreased relative to the full community (Figure S2AB). In contrast to the richness and diversity measurements, the Kullback–Leibler divergence compares the relative abundance of specific ASVs or OTUs between representations of the community. I calculated the Kullback–Leibler divergence between the full communities and those where rare ASVs were removed. As the threshold for removing ASVs increased, the amount of information lost also increased for both ASVs and OTUs (Figure 2C and Figure S2C). The relative loss of information was generally smaller for OTUs than than it was for ASVs. Removing rare ASVs, regardless of abundance threshold, had profound impacts on the representation of the communities.

**Figure 2.**
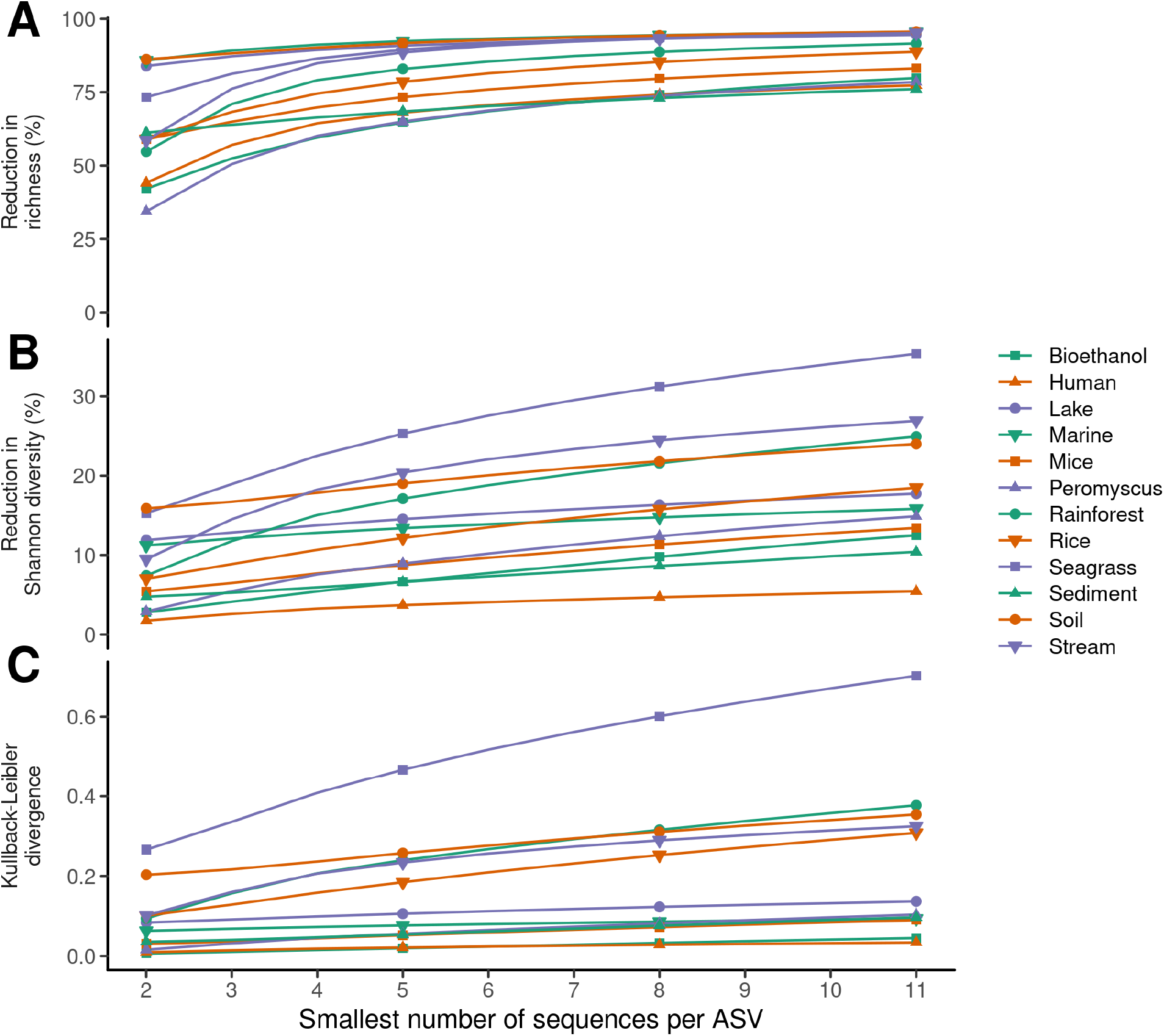
Removing rare sequences from samples alters their representation of alpha-diversity using amplicon sequence variants (ASVs). The average difference in the richness (A), Shannon diversity (B), and Kullback-Leiber divergence (C) for each sample within a dataset was calculated between the original community structures relative to applying different minimum abundance thresholds.

### Removing treatment group effects from community data

Because treatment effects often affect a sample’s diversity and inter-sample variation, I generated null distributions for each study by randomizing, without replacement, the number of times each ASV was observed in each sample such that the total number of sequences in each sample and the total number of times each ASV was observed across all samples in the study was the same as was originally observed. This effectively made every community in a study a statistical sample of the study-wide composite community distribution. For example, after this procedure, each of the 490 samples from the human dataset would be expected to have the same richness and diversity of ASVs and one would not expect to find treatment-based effects between the samples. Because of the risk of bias if only one representation of the null distribution was generated, I generated 100 randomized datasets for each study. The trends between removing rare ASVs and the richness, diversity, and information loss that were identified using the observed community distribution data were also identified with the data from the null distribution; however, the losses were larger when using the null distribution data (Figure S3). The null distribution data were used in the remainder of the study to minimize the risk of bias.

### The impact of removing rare ASVs on the information represented between samples

Considering the loss of richness, diversity, and information when a community has its rarest ASVs removed, it seemed likely that the relationship between communities would also be altered. To assess the impact of removing rare ASVs on measures of alpha diversity between samples I calculated the coefficients of variation (COVs, i.e. the standard deviation divided by the mean) for richness and diversity for each study at multiple abundance thresholds. The COVs for the richness of ASVs across the studies after removing singletons were between 3.6 and 32.7-times larger than they were without removing singleton ASVs (mice and stream; Figure 3A). Similarly, the COVs for the diversity of ASVs were between 1.8 and 20.4-times larger when singletons were removed than when they were not removed (mice and rice; Figure 3B). To assess the impact of removing rare ASVs on measures of beta diversity between samples, I calculated the COVs of the Bray-Curtis distances between samples within the same study at multiple abundance thresholds. The COVs between Bray-Curtis distances within a study when singletons were removed was between 1.3 and 18.6-times larger than when they were not removed (mice and stream; Figure 3C). When ASVs were clustered into OTUs the difference in COVs was less than it was for the ASVs (Figure S4). These results indicate that removing rare ASVs increases the dissimilarity between samples, which could have a significant impact on the statistical power to detect differences between treatment groups.

**Figure 3.**
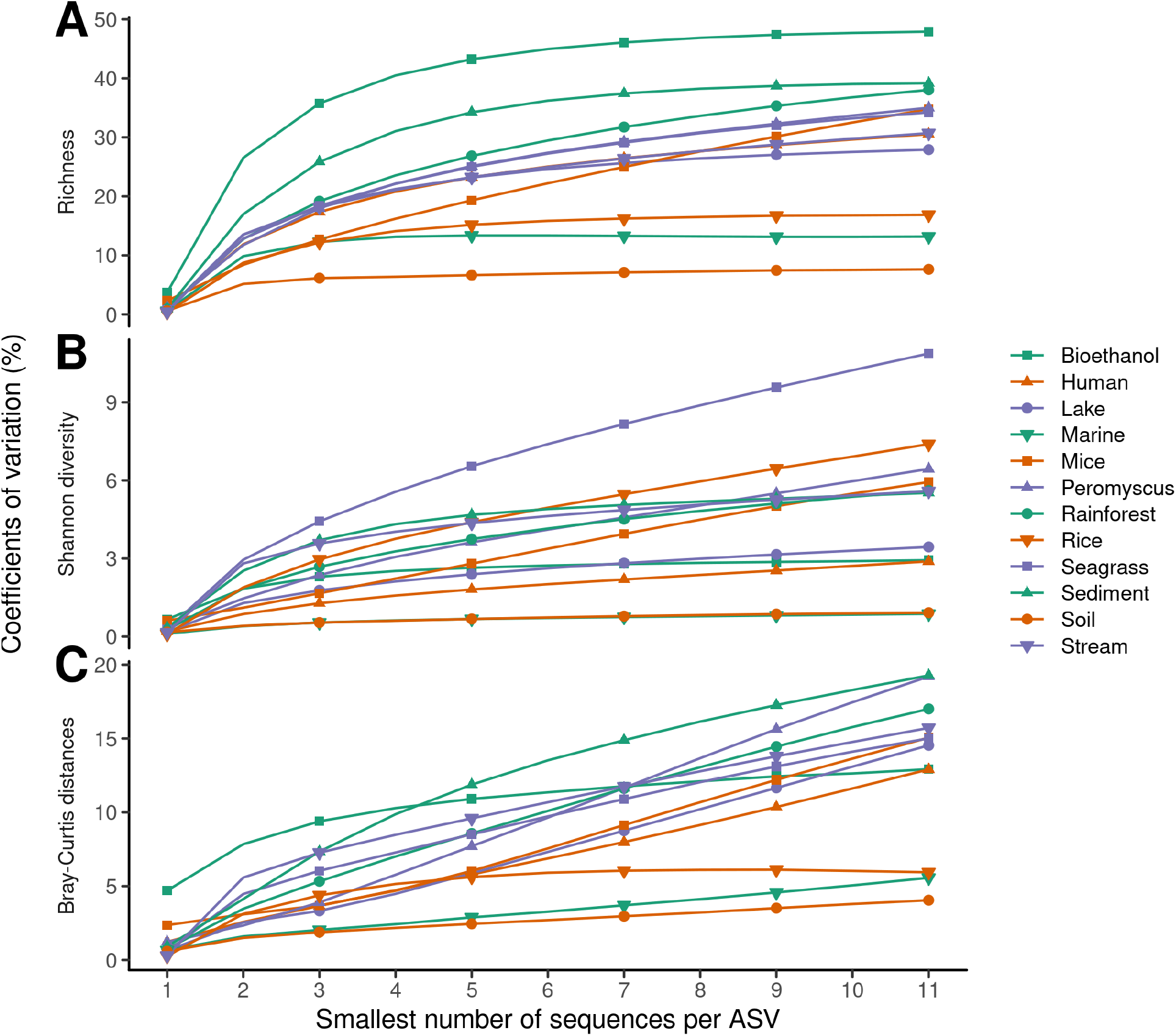
Removing rare sequences from samples increases the inter-sample variation for amplicon sequence variants (ASVs). The coefficient of variation in richness (A), Shannon diversity (B), and Bray-Curtis distances (C) for each dataset was calculated using the null distributed samples for each dataset with varying minimum abundance thresholds.

### The impact of removing rare ASVs on the ability to detect statistically significant differences between treatment groups

To test the effect of increased inter-sample variation, I randomly assigned samples to one of two treatment groups. In the first treatment group, communities were randomly sampled from the null distribution as described above. For the second treatment group, I randomly selected 10% of the ASVs in the pooled study distribution to increase their abundance by 5%. I randomly generated 100 simulated sets of treatment groups and samples. I then tested the ability to detect a difference between the two treatment groups using alpha and beta diversity metrics. The fraction of significant tests was a measurement of the statistical power to detect the difference between the treatment groups. When considering the differences in richness and diversity, the marine dataset yielded no simulated sets that were statistically significant, which was likely due to the small number of samples in the study (N=7). Among the remaining datasets, the power to detect a difference in the richness of ASVs ranged between 0.10 and 0.49 (sediment and stream) and between 0.10 and 0.53 (rainforest and stream) to detect a difference in diversity when using a Wilcox test (Figure 4A). When singleton ASVs were removed, the power to detect a difference in the richness of ASVs dropped by between 27.3 and 92.9% (bioethanol and soil) and by between 40.0 and 93.3% for their diversity(rainforest and soil; Figure 4B). The effect of removing rare ASVs on the richness of OTUs and their diversity was similar (Figure S5AB). I used the Bray-Curtis dissimilarity index to compare the simulated communities within each dataset and calculated the power to detect differences between the two simulated treatment groups using the analysis of molecular variance (Figure 4C and S5C). Without removing rare sequences, the power to detect a difference between the two simulated treatment groups varied between 0.41 and 1.00 (rainforest and rainforest). Aside from the bioethanol, human, and mice datasets, the power to detect differences dropped by between 6.5 and 64.0% (soil and rice) when singletons were removed. However, when ASVs that occurred 10 or fewer times were removed from each sample, the power to detect differences dropped by 12.0 and 97.2% (human and peromyscus); similar results were observed when ASVs were clustered into OTUs. Removing rare ASVs reduced the ability to detect simulated treatment effects using metrics commonly used to compare microbial communities.

**Figure 4.**
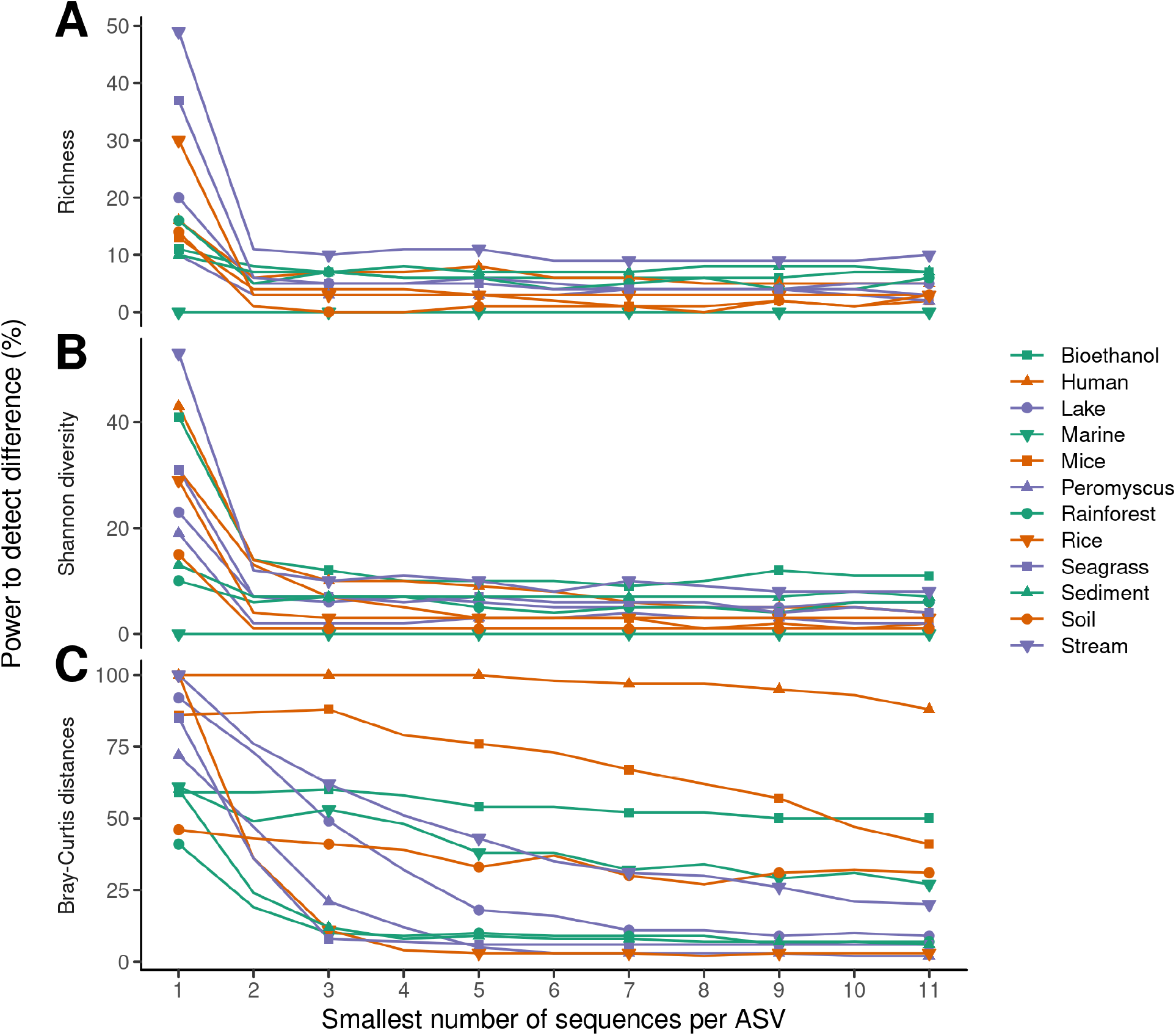
Removing rare sequences from samples reduces the statistical power to detect differences between empirically generated treatment groups when using amplicon sequence variants (ASVs). The fraction of significant tests comparing the richness (A) and Shannon diversity (B) using a Wilcox test and Bray-Curtis distances (C) using analysis of molecular variance for each dataset was calculated using empirically generated treatment groups containing equal numbers of samples for each dataset with varying minimum abundance thresholds. For each dataset and minimum abundance threshold, 100 randomizations were peformed.

### The impact of removing rare ASVs on the probability of falsely detecting a difference between treatment groups

I next asked whether removing rare ASVs could lead to falsely claiming that a treatment effect had a significant effect on community diversity and structure. First, I sampled sequences from the null distribution for each dataset and randomly assigned each sample to one of two treatment groups and determined the richness and diversity of ASVs and OTUs. Testing at an experiment-wise error rate of 0.05, I expected 5% of the iterations for each dataset to yield a significant test result. Indeed, there was no evidence that removing rare ASVs resulted in an inflated experiment-wise error rate. The average fraction of significant tests did not meaningfully vary from 0.05 across the minimum abundance threshold, dataset, metric of describing sample alpha-diversity, or whether the abundance of ASVs or OTUs were used (Figure 5A and S6A). Similarly, the average fraction of significant tests did not meaningfully vary from 0.05 when using analysis of molecular variance to compare communities using Bray-Curtis distances (Figure 5A and S6A). Second, I again sampled sequences from the null distribution, but assigned samples to one of two treatment groups based on the number of sequences in each sample. The samples with fewer than the median number of sequences for the dataset were assigned to one group and those with more than the median were assigned to the other. This exaggerated bias has been observed in comparisons of the lung and oral microbiota because of the larger number of non-specific amplicons that can be sequenced from lung samples relative to those in the oral cavity leading to a significant difference in sequencing depth between treatment groups (9). When rare sequences were not removed, the fraction of significant tests did not differ from 5% for comparing the richness, their diversity, or Bray-Curtis distances (Figure 5B and S6B). However, when rare taxa of any frequency were removed, the probability of falsely detecing a difference as signifiant increased with the definition of rarity (Figure 5B and S6B). Not including the small marine dataset, the average fraction the average fraction of falsely detecting a difference across datasets when only singletons were removed was 92.45%. If there is any relationship between the number of sequences and the treatment group, the risk of falsely rejecting the null hypothesis is inflated when researchers use the strategy of removing rare sequences. The most conservative approach is to not remove low abundance sequences.

**Figure 5.**
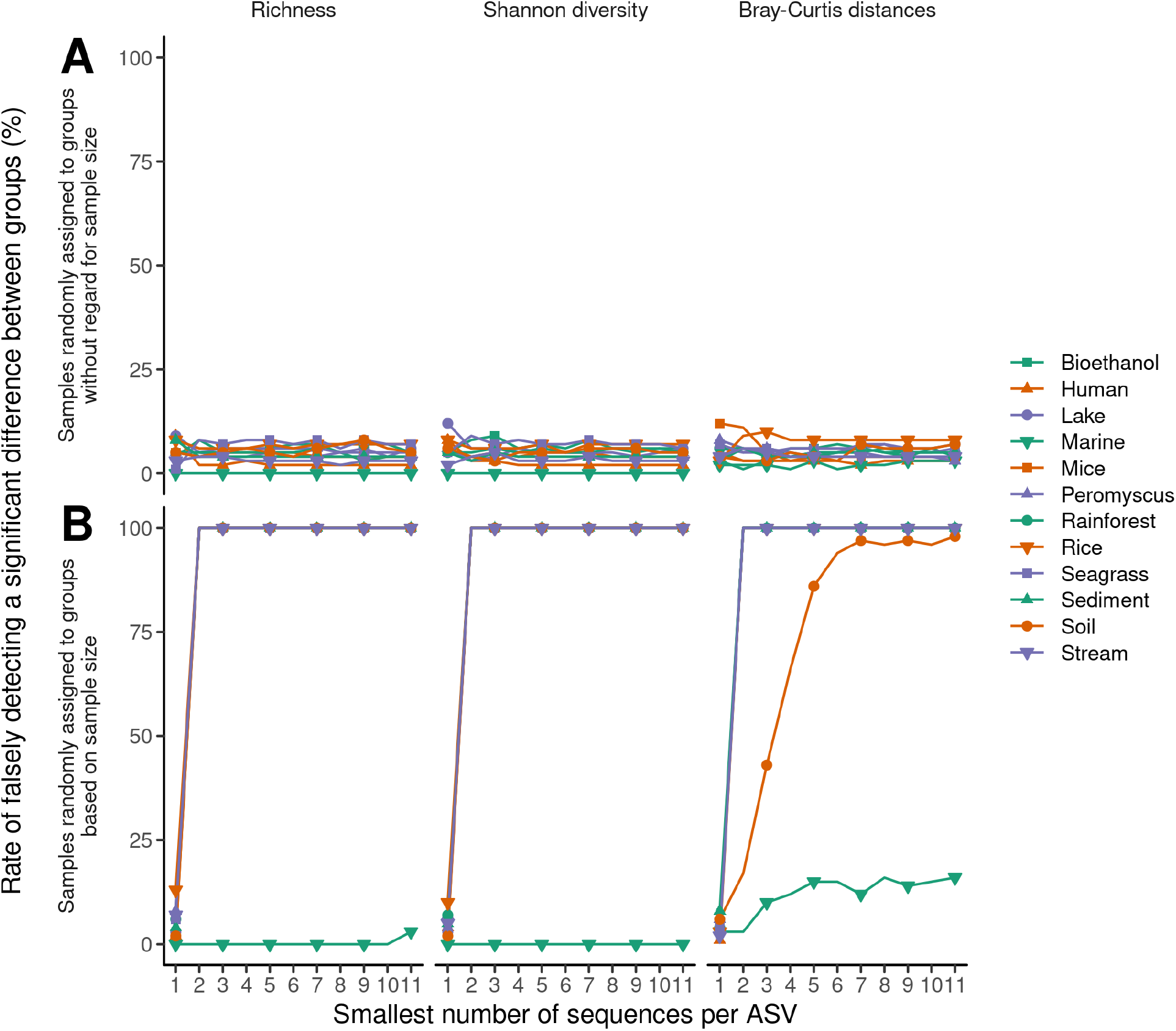
Removing rare sequences does not impact the false detection rate unless the number of sequences per sample is confounded with the treatment groups when using amplicon sequence variants (ASVs). The fraction of significant tests comparing the richness and Shannon diversity using a Wilcox test and Bray-Curtis distances using analysis of molecular variance for each dataset was calculated. Empirically generated treatment groups were generated containing equal numbers of samples where the samples represented a null distribution. In one simulation the samples were randomly assigned to a treatment group (A) and in the other the samples were assigned based on the number of sequences in each sample (B). For each dataset and minimum abundance threshold, 100 randomizations were peformed.

## Discussion

Removing rare sequences from 16S rRNA gene sequence data is a common practice that is used as a heuristic to help remove residual PCR artifacts and low quality sequences. In this analysis, I have shown that rare sequences are more common in samples with shallow sequencing than in those with deep sequencing and that rare sequences are frequently observed in multiple samples. These observations suggest that many of the sequences being removed are actually good sequences. Becuase rarity is often defined by a fixed number of observations per sample (e.g. sequences that only appear once in a sample, regardless of the size of the sample), removing rare sequences has a disproportionate impact on samples with fewer sequences. Removing rare sequences resulted in a reduction in the alpha diversity and a pronounced change in the structure of individual samples. The effect was an increase in the differences observed between samples, which made it more difficult to detect differences between treatment groups when differences actually existed. Furthermore, if the number of reads per sample was confounded with the treatment groups, then removing rare sequences increased the probability of falsely detecting a difference between the treatment groups. The practice of removing rare sequences from samples should be stopped.

The practice of removing rare sequences from samples seems to be a response to researchers prioritizing the number of reads and length of sequences over their quality (5). Previous work has shown that assembly of fully overlapping sequence reads results in the lowest sequencing error rates (2). The studies highlighted in this analysis sequenced the V4 region of the 16S rRNA gene using the Illumina MiSeq sequencing platform with their version 2 chemistry. The resulting data consists of two 250 nt reads that span a region that is about 253 nt long. In contrast, there has been a movement to sequence longer regions with similar chemistry resulting in less overlap between the sequencing reads (10, 11). Alternatively, others have prioritized increasing the number of sequences per sample by sequencing the V4 region, but with paired or single reads that are 150 nt long (12, 13). Both practices result in a significantly higher error rate for the resulting assmebly. Instead, researchers should prioritize the quality over the quantity and length of their data. For these reasons, this analysis did not use lower quality data generated by alternative methods.

The impacts of removing rare sequences on the representation of communities are caused by the uneven impact of applying a single abundance threshold across all samples. The number of sequences per sample in the datasets highlighted in this analysis varied by as much as 100-fold (Table 1; Figure S1). In practice, I suspect this range is actually larger since researchers may have opted against depositing samples with fewer reads into the SRA. If the range in read coverage for a study is 100-fold, a singleton in a small sample would have had a comparable relative abundance as a sequence that appeared 100 times in a more densely sequenced sample. However, it would have been removed from the smaller sample and not the larger sample. Thus, applying a single abundance threshold disproportionately impacted the samples with fewer sequences. This seems to contradict the purpose of removing rare sequences since a singleton in the smaller sample is more reliable than the singleton in the larger sample. A superior approach to removing rare sequences is to use rarefaction to conrol for uneven sampling.

Even with the best sequencing approaches, PCR artifacts and sequencing errors persist. This may account for some sequences not being observed other samples. Although this lack of inter-sample coverage could have been due to treatment effects and natural variation, it is likely that some portion of the sequences that were unique to samples were artifacts or contained errors. Researchers are encouraged to use rarefaction to control for uneven sampling and the presence of spurious sequences. Previous work sequencing mock communities has shown that the number of spurious sequences increases with sequencing depth (2, 14). By rarefying data to a common number of sequences per sample, the number of spurious sequences can be controlled. As shown in the data I presented, which was rarefied to a common number of reads per sample within a dataset, when rare sequences were not removed the power to detect differences was the highest and the false discovery rate was the expected 5% (Figures 5 and S6).

In addition to considerations of how to control for the presence of spurious sequences, researchers also need to be mindful of how to interpret the results of their work. Because every dataset will contain residual sequences that are spurious, measures of richness and diversity should be made on a relative basis. For example, pronouncements that communities from an environment or treatment contain a specific number of taxa are problematic. Instead, we should limit ourselves to indicating that samples from one treatment group has more taxa than another without using absolute values of richness. Furthermore, we must take caution in interpreting rare taxa. Although the data from the studies highlighted here suggest that most rare sequences are not spurious, it is likely that some are. Therefore, researchers must approach rare sequences with more skepticism than more abundant sequences. Reseachers should seek out other methods to confirm inferences that they make about rare sequences. This is a standard that should be applied regardless of their abundance (15).

How to curate and interpret rare sequences has been a significant challenge since microbial ecologists transitioned away from Sanger sequencing of samples (16–18). Although the extent of the “rare biosphere” is still an open question, it is important to appreciate the importance of rare populations in all communities. Populations can be numerically rare but ubuiquitous or abundant and limited in their geographic range. Alternatively, they can be numerically rare but temporally common or abundant but present infrequently. Removing sequences from any of these settings will limit our ability to study the role of such populations or the processes that drive their patchy distributions that are so common with microbial communities (8).

## Materials and Methods

### Data curation and analysis

To insure the highest possible data quality, datasets were limited to those where the 500 cycle version 2 MiSeq chemistry was used to sequence the amplicons. The paired 250 nt reads resulted in near complete 2-fold sequencing coverage of every nucleotide in the ca. 253 nt-long region. This region and sequencing platform were selected because previous work has shown that a standard data analysis pipeline in mothur results in a sequencing error rate below 0.02% (2). All sequence data were obtained from the Sequence Read Archive (SRA) and processed using a standard mothur-based sequencing pipeline that resulted in ASVs as generated by the pre.cluster algorithm using a threshold of 2 nt (2, 19). ASVs were assigned to OTUs using a 3% distance threshold using mothur’s cluster function with the OptiClust algorithm (20). To minimize the effects of uneven sampling effort, samples were rarefied to the number of sequences in the smallest sample for each dataset. Because metrics of alpha diversity did not consistently follow a normal distribution, I used the non-parametric Wilcoxon rank test as implemented in R. For comparisons of Bray-Curtis distances, the amova function within mothur was used, which implements the analysis of molecular variance algorithm (21).

### Reproducibility

All analyses were performed using mothur (version 1.44.1) and R (version 4.0.2) with the tidyverse (version 1.3.0), broom (version 0.7.0), data.table (version 1.13.4), and cowplot (version 1.1.0) packages. All of the code that was used in this analysis as well as the Makefile for running the analysis are available at https://github.com/SchlossLab/Schloss_Singletons_XXXXX_2019 along with the complete version history of the project.

## Acknowledgements

I endebted to the researchers who developed the 12 datasets used in this study for depositing their sequence data into the Sequence Read Archive. This work was supported in part by funding from the National Institutes of Health (U01AI124255, P30DK034933, R01CA215574).

---

**Figure S1.**
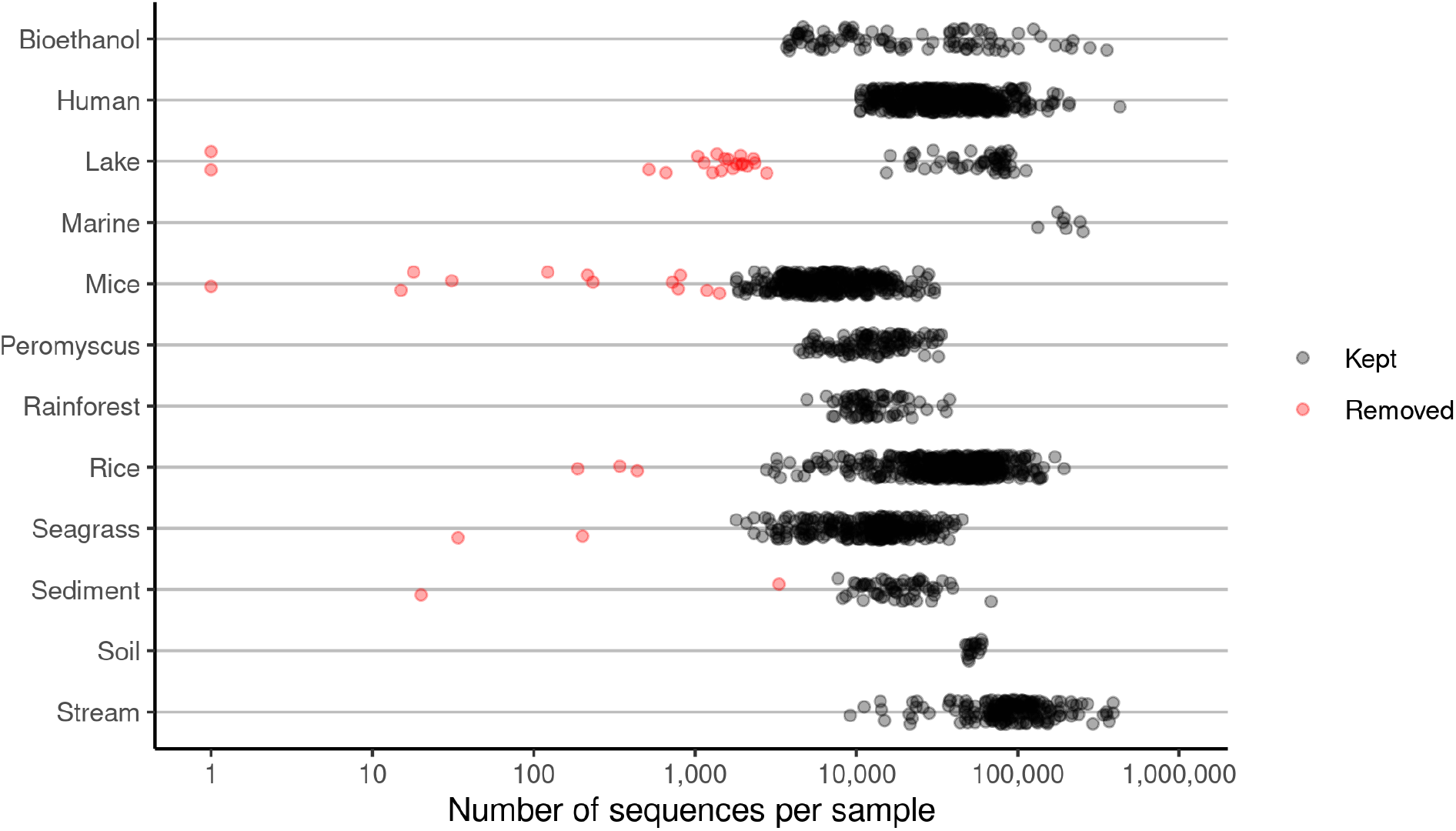
Distribution of the number of sequences per sample in the 12 datasets included in this study. A different minimum number of sequences per sample threshold was applied to each dataset based on identifying natural breaks in the distribution of the number of sequences per sample.

**Figure S2.**
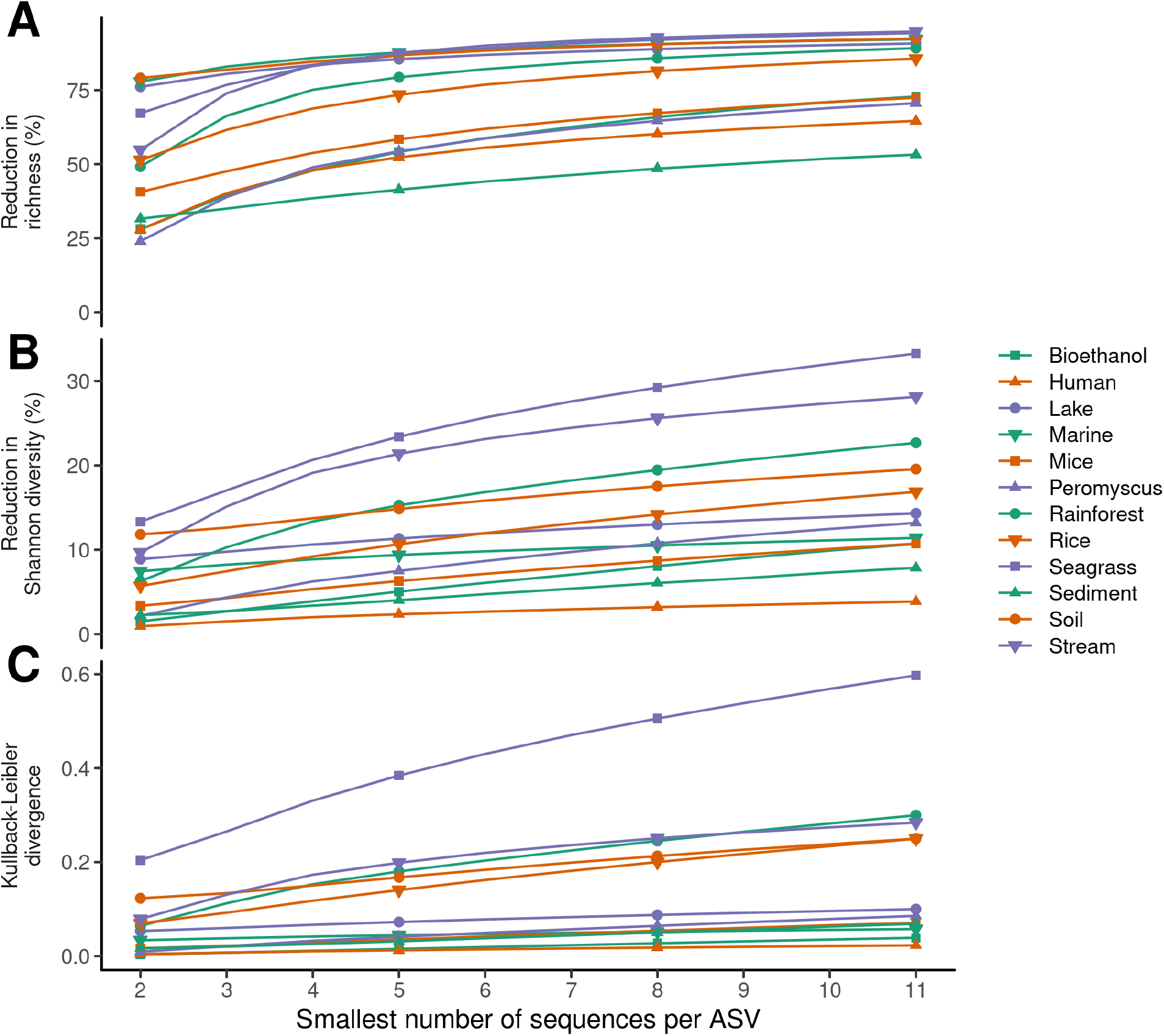
Removing rare sequences from samples alters their representation of alpha-diversity using operational taxonomic units (OTUs). The average difference in the richness (A), Shannon diversity (B), and Kullback-Leiber divergence (C) for each sample within a dataset was calculated between the original community structures relative to applying different minimum abundance thresholds.

**Figure S3.**
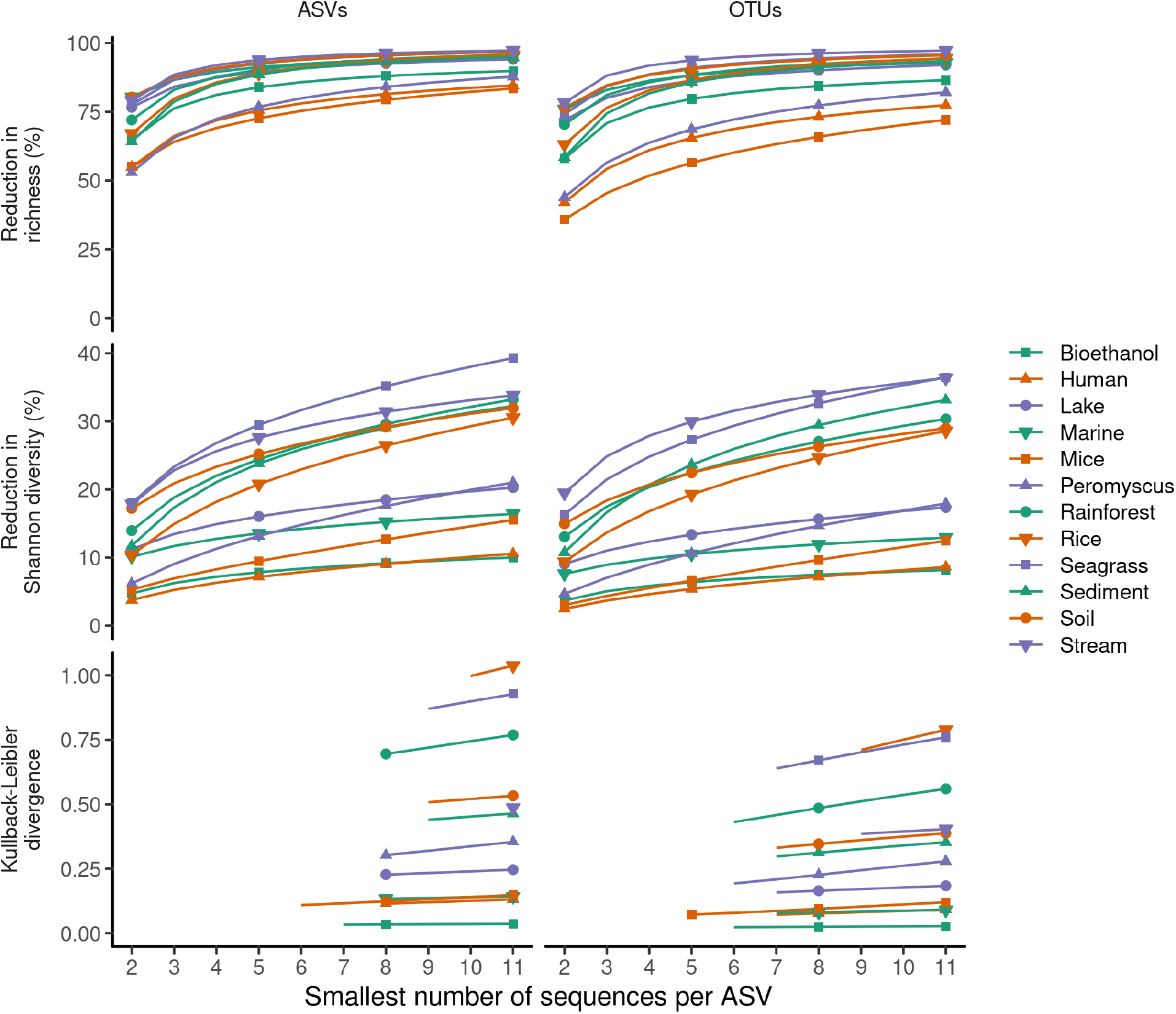
Removing rare sequences from samples alters their representation of alpha-diversity when regenerating samples using a null distribution for each dataset. The average difference in the richness, Shannon diversity, and Kullback-Leiber divergence for each sample within a dataset was calculated between the original community structures relative to applying different minimum abundance thresholds. Some values of Kullback-Leiber divergence are missing because undefined values were calculated due to the removal of rare sequences. Data are shown for amplicon sequence variants (ASVs) and operational taxonomic units (OTUs)

**Figure S4.**
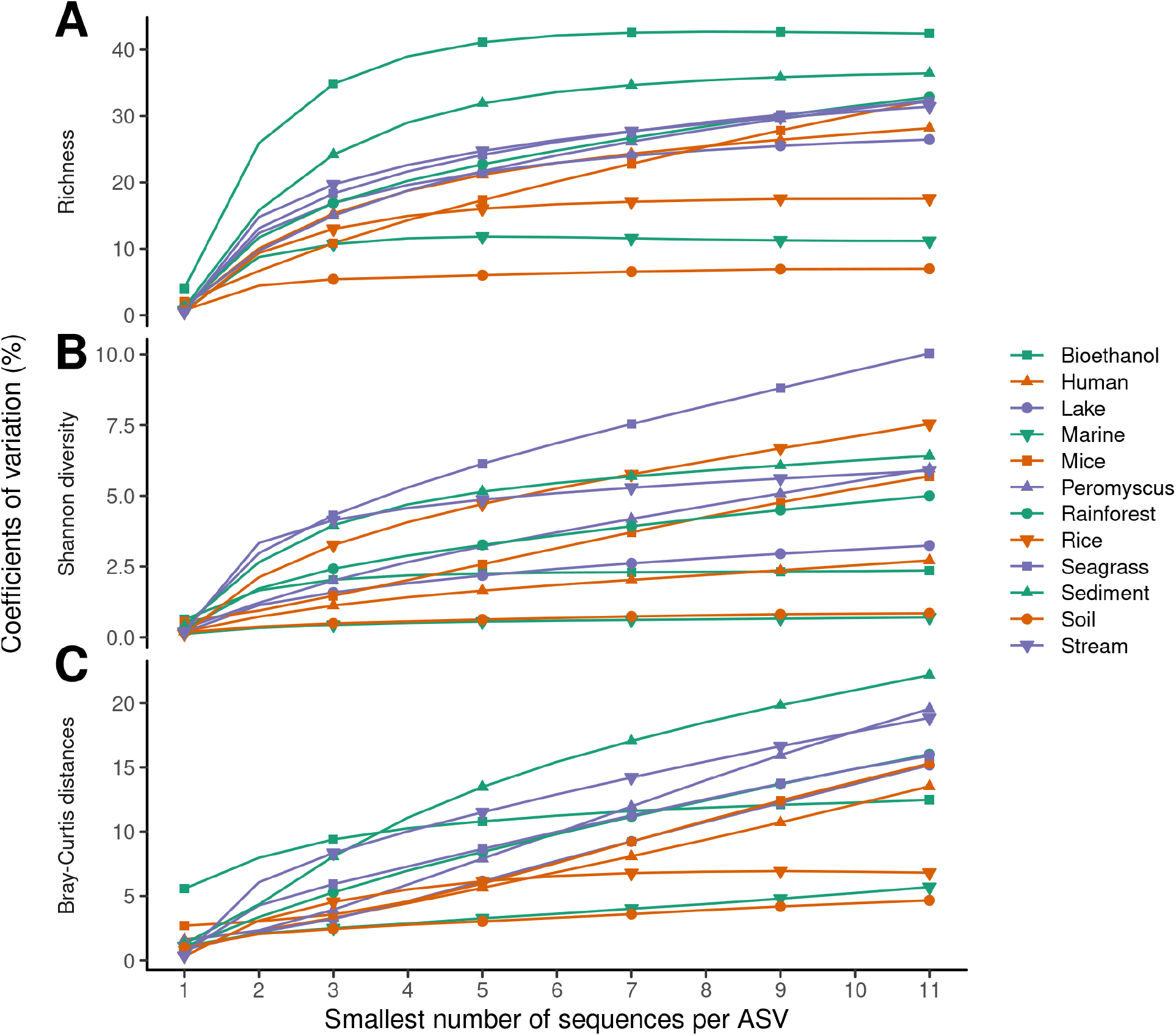
Removing rare sequences from samples increases the inter-sample variation for operational taxonomic units (OTUs). The coefficient of variation in richness (A), Shannon diversity (B), and Bray-Curtis distances (C) for each dataset was calculated using the null distributed samples for each dataset with varying minimum abundance thresholds.

**Figure S5.**
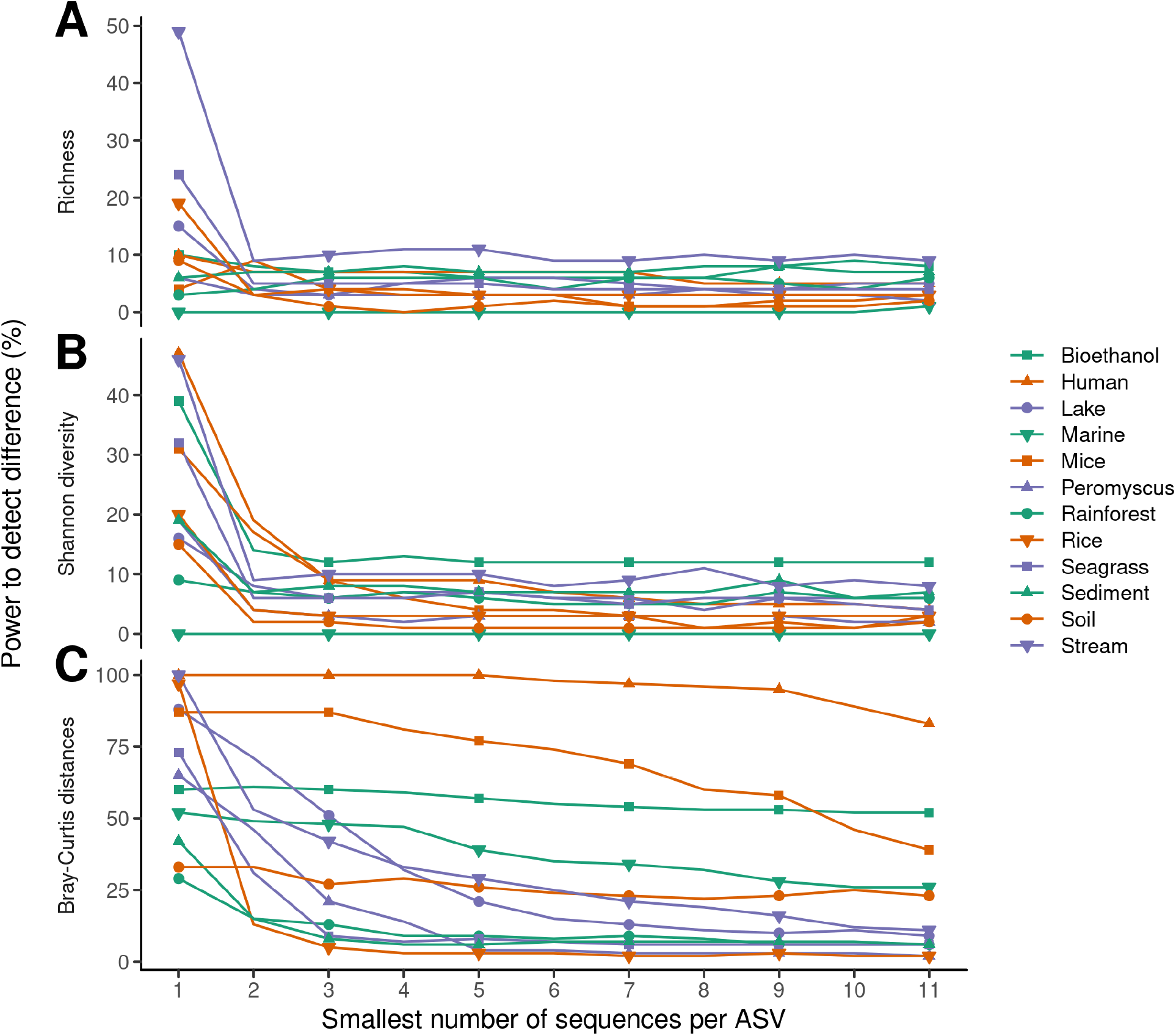
Removing rare sequences from samples reduces the statistical power to detect differences between empirically generated treatment groups when using operational taxonomic units (OTUs). The fraction of significant tests comparing the richness (A) and Shannon diversity (B) using a Wilcox test and Bray-Curtis distances (C) using analysis of molecular variance for each dataset was calculated using empirically generated treatment groups containing equal numbers of samples for each dataset with varying minimum abundance thresholds. For each dataset and minimum abundance threshold, 100 randomizations were peformed.

**Figure S6.**
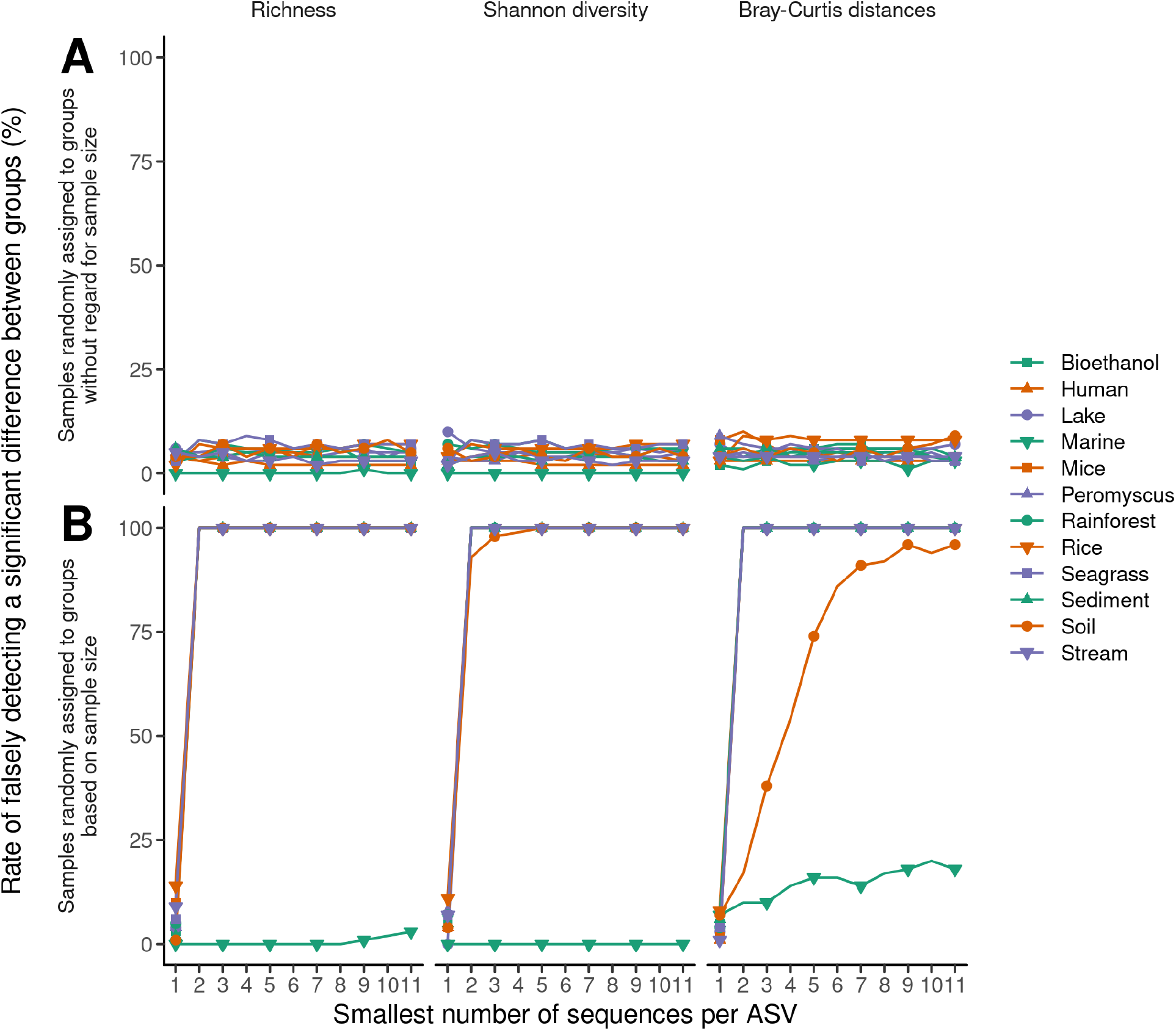
Removing rare sequences does not impact the false detection rate unless the number of sequences per sample is confounded with the treatment groups when using operational taxonomic units (OTUs). The fraction of significant tests comparing the richness and Shannon diversity using a Wilcox test and Bray-Curtis distances using analysis of molecular variance for each dataset was calculated. Empirically generated treatment groups were generated containing equal numbers of samples where the samples represented a null distribution. In one simulation the samples were randomly assigned to a treatment group (A) and in the other the samples were assigned based on the number of sequences in each sample (B). For each dataset and minimum abundance threshold, 100 randomizations were peformed.

